# High-Throughput Functional Assay in Cystic Fibrosis Patient-Derived Organoids Allows Drug Repurposing

**DOI:** 10.1101/2022.07.14.500147

**Authors:** S. Spelier, E. de Poel, G.N. Ithakisiou, S.W.F. Suen, M.C. Hagemeijer, D. Muilwijk, A.M. Vonk, J.E. Brunsveld, E. Kruisselbrink, C.K. van der Ent, J.M. Beekman

**Author notes:** These authors contributed equally (S. Spelier, E. de Poel). **TAKE-HOME MESSAGE** We established a high-throughput functional assay using CF patient-derived intestinal organoids. We screened 1400 FDA-approved compounds and found that statins increased function of W1282X/W1282X CFTR, when combined with CFTR modulators.

## Abstract

Cystic fibrosis (CF) is a rare hereditary disease caused by mutations in the CFTR gene. Recent therapies enable effective restoration of CFTR function of the most common F508del CFTR mutation. This shifts the unmet clinical need towards people with rare CFTR mutations such as nonsense mutations, of which G542X and W1282X are most prevalent. CFTR function measurements in patient-derived cell-based assays played a critical role in preclinical drug development for CF and may play an important role to identify new drugs for people with rare CFTR mutations.

Here, we miniaturized the previously described forskolin induced swelling (FIS) assay in intestinal organoids from a 96-wells to a 384-wells plate screening format. Using this novel assay, we tested CFTR increasing potential of a 1400-compound FDA-approved drug library in organoids from donors with W1282X/W1282X CFTR nonsense mutations.

The 384-wells FIS-assay demonstrated uniformity and robustness based on CV and Z’-factor calculations. In the primary screen, the top 5 compound combinations that increased CFTR function all contained at least one statin. In the secondary screen, we indeed verified that four out of the five statins, Mevastatin; Lovastatin; Simvastatin and Fluvastatin increased CFTR function when combined with CFTR modulators. Statin-induced CFTR rescue was W1282X specific, as increased CFTR function was not shown for patient-derived organoids harbouring R334W/R334W and F508del/F508del mutations.

Future studies should focus on elucidating genotype specificity and mode-of-action of statins into more detail. This study exemplifies proof-of-principle of large-scale compound screening in a functional assay using patient derived organoids.

**Graphical abstract:** 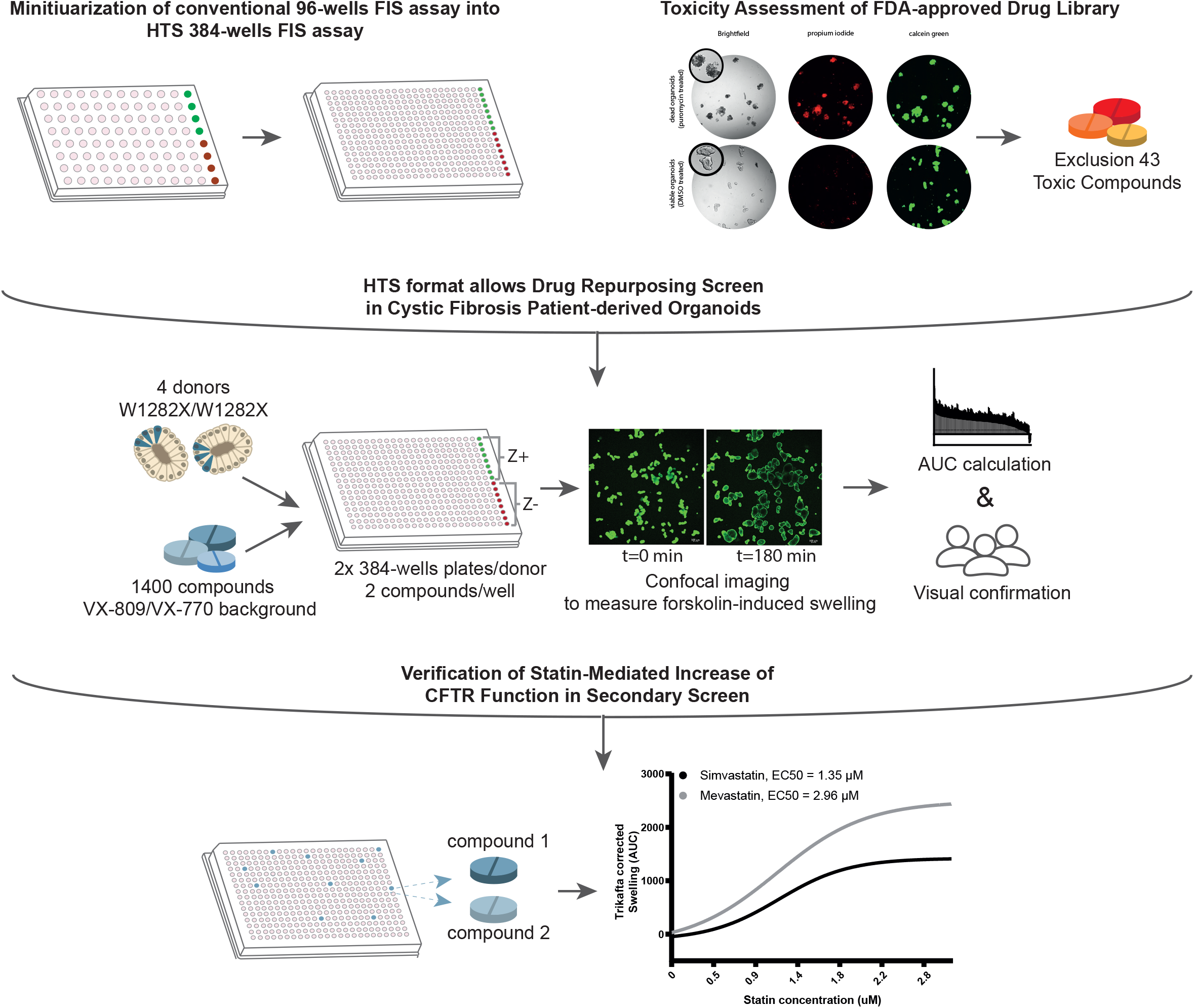

## INTRODUCTION

Cystic fibrosis (CF) is a rare, monogenic disease that is caused by mutations in the cystic fibrosis transmembrane conductance regulator (CFTR) gene. Mutations in CFTR lead to loss of chloride secretion and deficient fluid transport across tissue epithelium. Subsequently, thick and sticky mucus secretions are produced resulting in chronic bacterial infections, progressive loss of pulmonary function and multiorgan failure ^1^. More than 1700 distinct mutations have been estimated to be disease-causing ^2^. The F508del mutation is mainly prevalent, occurring on approximately one allele in 85% of the CF patients. For those patients, increasingly effective treatments called CFTR modulators have been developed in the last decade aiding in proper CFTR protein folding and membrane potentiation ^3^. Recently the potent combination of the three CFTR modulators VX-445/VX-661/VX-770 has been approved by the FDA and EMA. However, an urgent need for effective restoration of CFTR function persists for the remaining CF patients that cannot benefit from CFTR modulation.

Approximately 10% of the worldwide CF population carry premature termination codon (PTC) mutations, resulting in production of truncated CFTR protein. Previous preclinical research identified compounds with read-through (RT) activity that introduce an amino acid at the PTC site and thereby results in full-length protein production ^4,5^. Clinical efficacy of RT approaches is however limited, as demonstrated by for example clinical studies with RT agent Ataluren/PTC-124 ^6^. A novel recently developed RT agent ELX-02 (NB124; Eloxx Pharmaceuticals) resulted in production of full length CFTR protein and restoration of CFTR function in G542X/G542X intestinal organoids ^7^. However, in a study characterizing the effect of ELX-02 in a larger set of patient-derived organoids, ELX-02 as single agent resulted in only limited restoration of CFTR function ^8^. In a recent press release by Eloxx Pharmaceuticals, first results of their phase 2 trial in homozygous G542X CF patients describe only a minor decrease in sweat chloride levels (‘Eloxx Pharmaceuticals Reports’, 2021). Altogether, this underlines the need for continuing the search for novel CFTR modulating molecular entities for people with CFTR PTC mutations.

Drug development in the context of rare diseases where the numbers of patients are low is challenging. For these patient populations, repurposing of clinically approved drugs could be beneficial. Drug repurposing is an attractive approach that aims to extend the indication of already marketed drugs ^10^. A prerequisite for drug repurposing is that the exploited assay is robust and associates with clinical features of disease, such as therapeutic response. CFTR function measurements in patient-derived intestinal organoids (PDIOs) associate with clinical features of CF and may enable drug repurposing in a personalized setting. We previously developed a PDIO-based assay for CF based on forskolin-induced swelling (FIS). Forskolin (fsk) induces fluid secretion into the organoid lumen resulting in CFTR-dependent swelling, which can be quantified with automated live-cell-microscopy ^11,12^. CFTR function measurements in PDIOs associate with (a) disease severity indicators of CF, (b) long-term disease progression and (c) therapeutic response, enabling preclinical drug efficacy and mode-of-action studies ^13,14^. As such, the FIS assay is well-suited for identifying and prioritizing drugs that influence CFTR function. However, the 96-wells format of the assay limits its use when higher throughput screening is required.

Here, we miniaturized the 96-wells FIS assay towards a 384 wells-plate format enabling drug screening in PDIOs at higher throughput. Important factors when validating assay quality are within and between experiment variability, drift and edge-effects within plates and comparison of the outcomes of experiments performed with the original 96-wells set-up to the novel 384-wells set-up ^15^. As these validations yielded positive results, Z’-factors were calculated to ultimately summarize assay performance ^16^. The 384-wells FIS assay was subsequently used to identify compounds from an FDA-approved, commercially available drug library for their capacity to increase CFTR function. The screen was performed in PDIOs harboring PTC W1282X/W1282X CFTR, as W1282X encompasses 18% of all PTC mutations, thereby being the most common PTC mutation in CF patients after G542X (CFTR2 database, 2021). Additionally, previous studies have shown that W1282X CFTR is to a limited extent responsive to CFTR modulators such as VX-445/VX-661/VX-770 ^18^. As such, exploiting PDIOs with this genotype allows potential identification of hits that can improve organoid swelling through various mode-of-actions, ranging from nonsense-specific effect to more general CFTR modulating effects.

In brief, we show proof-of-principle of a HTS assay using PDIOs in a functional assay. By screening a library of FDA-approved compounds, we pave the way for drug repurposing for people with CF for whom no clinical options are available.

## Results

### The 384-wells FIS-assay is reproducible, spatial uniform and has a comparable dynamic range to the 96-wells FIS-assay

First, we adapted several practical aspects of the 96-wells FIS format which was previously described by Vonk et al ^12^ to allow a higher-throughput working method (**Table 1**). In brief, 384-well plates were pre-cooled prior to organoid addition, a higher volume of a lower matrigel-percentage was added to each well to cover the whole well-surface and during image acquisition X/Y/Z locations were based on autofocus.

**Table 1.** Adaptations in FIS protocol of original 96-wells format to allow a 384-wells format

|  | 96-wells format, as described by<sup>12</sup> | 384-wells format |
| --- | --- | --- |
| 1 | Plates are prewarmed at 37°C prior to organoid-matrigel addition | Plates are precooled at -20°C, and kept on ice during organoid-matrigel plating |
| 2 | 4 µl 50% matrigel organoid suspension is added per well as drops | 10 µl 25% matrigel organoid suspension is added per well with an automatic multichannel, to cover the whole surface |
| 3 | - | Plate is spun down in a centrifuge to ensure matrigel coverage of the whole well surface |
| 4 | After 10 minutes at 37°C, 50 µl medium per well is added | After 10 minutes at 37°C, 8 µl medium per well is added |
| 5 | X/Y/Z location is manually set for each well | X/Y/Z is automatically set based on plate lay-out and autofocus of 12 reference points |

Using these adjustments, we verified the quality and reliability of the 384-wells FIS assay. In **Fig. 1A** we summarize the advised^15^ steps of assay quality validation when miniaturizing a cell-based assay. To demonstrate FIS assay reproducibility, we performed replicate experiments in two organoid lines with different genotypes. FIS was assessed in F508del/S1251N organoids treated with CFTR potentiator VX-770 and in F508del/F508del organoids treated with CFTR potentiator VX-770 and CFTR corrector VX-809. Minimal (min) and maximal (max) signal of swelling was induced with 0.008 and 5 µM fsk, respectively. The mean as well as the spread of the min and max signals were comparable between the replicate experiments both for F508del/S1251N organoids (**Fig. 1B & 1C)** and F508del/F508del organoids (**Fig. 1D & 1E**). In order to further characterize precision and repeatability within and between these replicate experiments, coefficient of variation (CV) were calculated ^19^. For the max signal, the average intra-assay CV values were 16% and 13% for the F508del/F508del and F508del/S1251N organoids respectively, the inter-assay CV values were respectively 4% and 8%. CV values should not exceed 20% and CV values under 10% are considered excellent, underlining the reproducibility of the 384-wells FIS assay ^15^.

**Figure 1:**
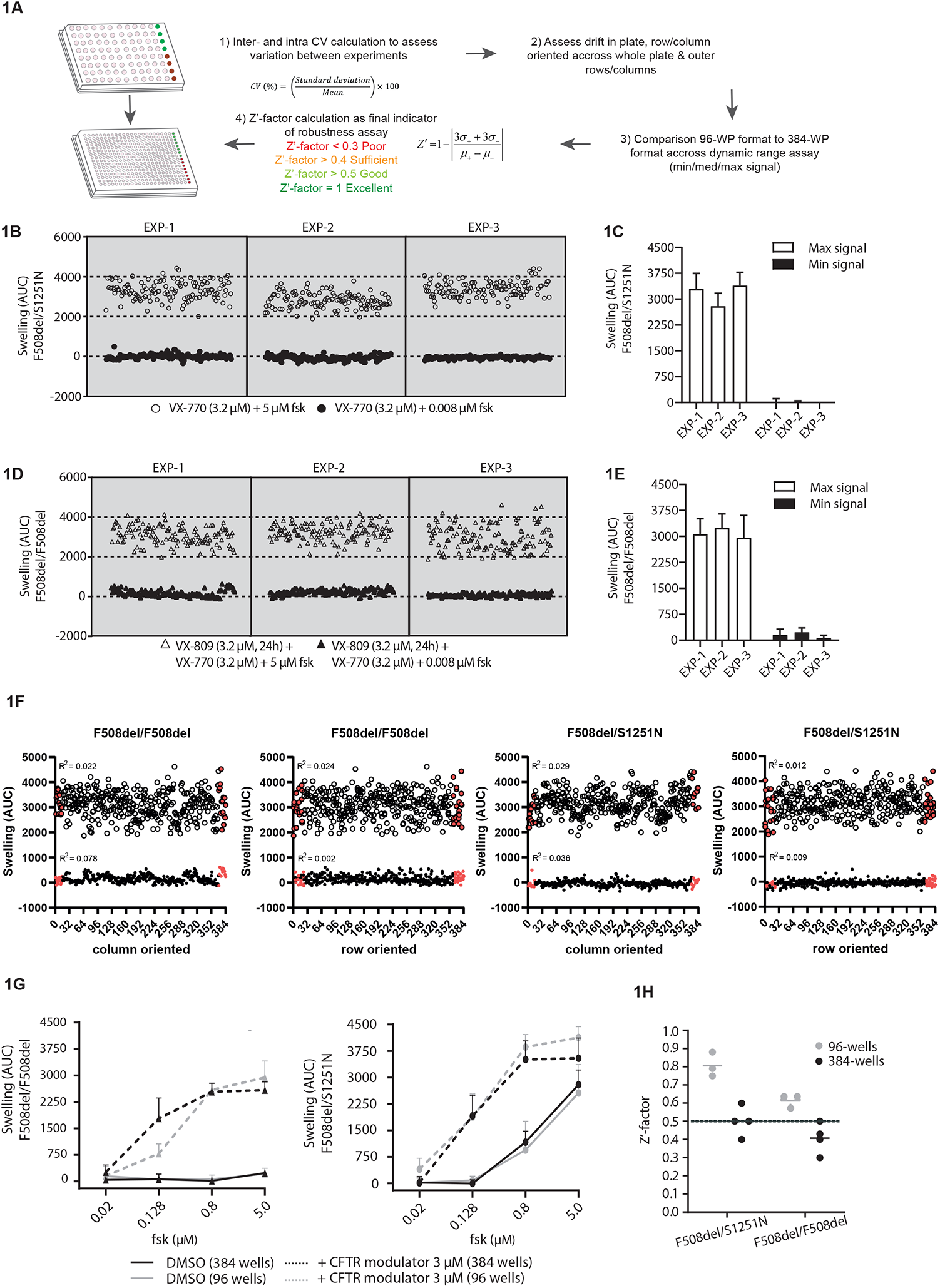
Reproducibility and dynamic range of 384-wells FIS assay. (A) Summary of optimization steps for assay development, in this paper specifically for miniaturizing a cell-based assay from 96-wells format to 384-wells format.(B) & (D) Replicate experiments of three 384-wells plates with respectively F508del/S1251N and F508del/F508del PDIOs, performed on three different culturing days. VX-770 (3.2 µM) and fsk (0.008 µM) was added to 128 wells of each 384-well plate to induce minimal swelling (=min signal wells). Maximum swelling was achieved by the addition of 5 µM fsk and VX-770 (3.2 µM) to another 128 wells (=max signal wells). The mean+SD swelling of all min and max signal wells of each plate depicted in B and D are summarized in (C) and (E) respectively. (F) Replicate experiment of three 384-wells plates with F508del/F508del and F508del/S1251N PDIOs redepicted separately in a row-oriented and column oriented fashion, to explore horizontal or vertical drift. For both the high and the low swelling condition, R^2^ was calculated. Outer rows and columns are depicted as red dots. (G) Swelling of F508del/F508del PDIOs treated with VX-770/VX-809 and F508del/S1251N PDIOs treated with VX-770 and in presence of an increasing concentration of fsk, measured in 96-wells plates and 384-wells plates (datapoints represent mean+SD, n=3 for the 96-wells experiments, n=1 for the 384-wells experiments). (H) Z’-factors calculated of each replicate experiment plate included in (C), (E) and (G).

Next, edge effect for both outer rows and columns as well as overall horizontal or vertical drift on the plates was characterized (**Fig. 1F**). As R^2^ values reach 0 in both donors for both the row and the column-oriented data, no horizontal or vertical drift across the plates is present. Additionally, the max AUC values of the outer rows and columns (indicated with red dots) do not differ significantly from the rest of the data in both donors.

We next compared the signal amplitude of the 384-wells FIS-assay with the conventional 96-wells assay. F508del/F508del and F508del/S1251N organoids with or without CFTR modulator(s) were stimulated with an increasing concentration of fsk (**Fig 1G**). This allows characterization of “medium” swelling levels in the FIS assay, which is crucial when characterizing the capacity of the assay to capture hit compounds during a screen (Chai, 2015). Medium signal needs to lie between the negative and positive controls and we show that indeed at 0.128 and at 0.8 µM fsk the mid-range of the assay is covered. Differences between the 96-wells and 384-wells assay are negligible, except for 0.128 µM fsk in the F508del/F508del organoids. Lastly, Z’-factors were calculated as indicator of assay quality. The Z’-factor is a parameter based on positive and negative control that ranges between 0 and 1, with 1 indicating a perfect assay and Z’-factors larger than 0.4 considered acceptable (Chai, 2015). Whilst mean Z’-factors from the 96-wells format experiments were higher compared to those of the 384-wells format experiments (**Fig. 1H**), the mean Z’-factor of the F508del/S1251N organoids was 0.4 and the mean Z’-factor of the F508del/F508del organoids was 0.5, indicating adequate assay robustness.

### Organoid viability is affected by 43 out of 1443 FDA-approved compounds

Prior to assessing the effect of compounds on CFTR function restoration, we assessed potential toxicity of the 1443 FDA-approved compounds. Organoid viability was determined by a dual live cell staining approach in which calcein was used to stain metabolically active cells and propidium iodide (PI) to visualize dead cells, a toxicity assay that has previously been performed on intestinal organoids ^20,21^. The ratio of total calcein green area and total PI area was calculated to correct for varying number of organoids between wells, after which values were normalized to Z-scores facilitating comparisons between plates **(Fig. 2A)**. Z-scores beyond 2, indicating toxicity, were found for 41 compounds (**Fig. 2B**). Additionally, organoid morphology was verified by three blinded observers. This led to a further exclusion of two additional compounds. The majority of the toxic compounds (83%), listed in **Supplemental Table 1**, are described as anti-cancer drugs. All 43 compounds were excluded from further screening.

**Figure 2:**
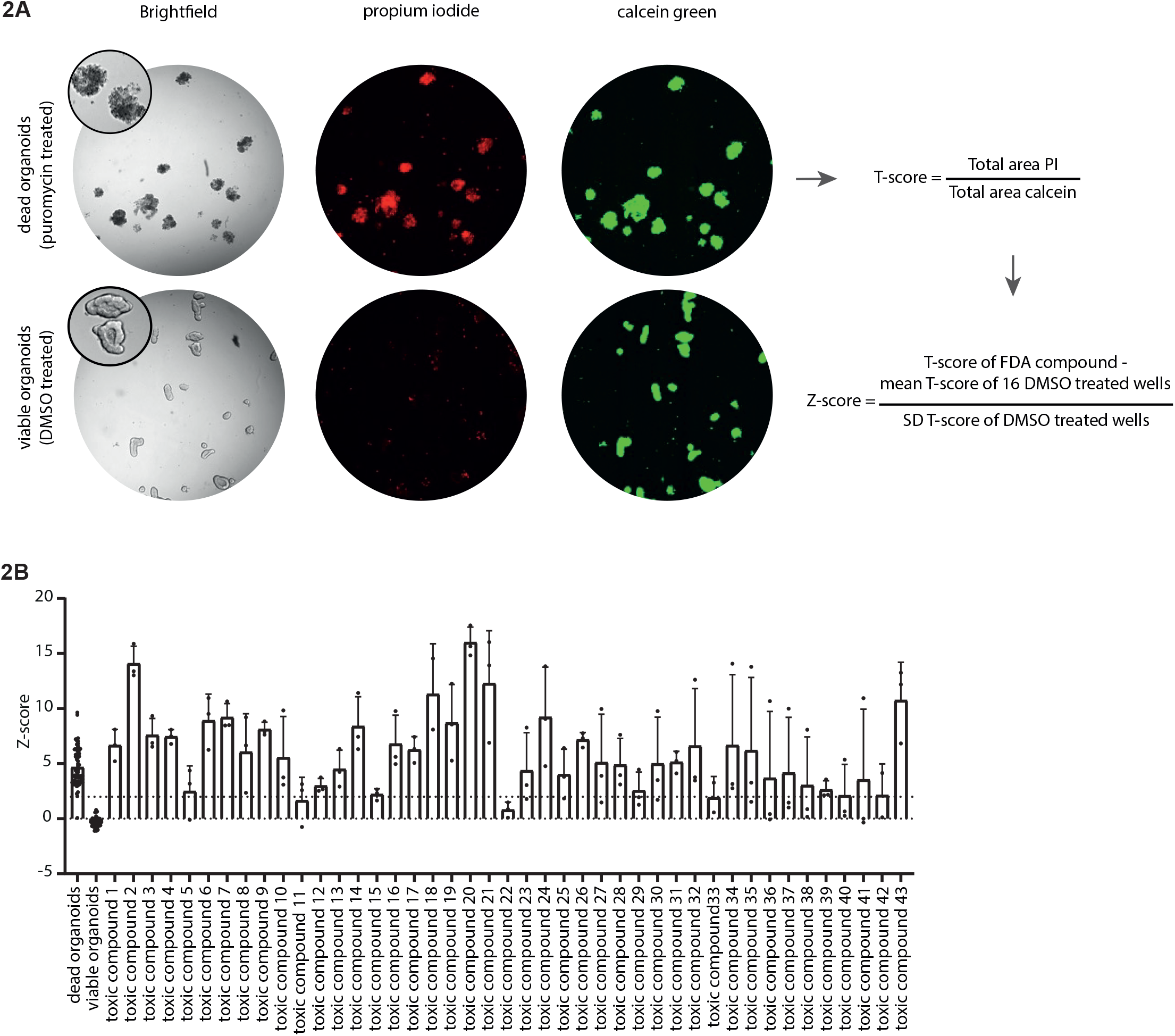
Organoid viability upon treatment with 1443 FDA-approved drugs. (A) Schematic of toxicity assessment pipeline. The ratio of total area calcein green and total area PI (T-score) was calculated to correct for varying number of organoids between wells. To further normalize to the negative control organoids, the Z-score was calculated between the compound-treated organoids and control (DMSO-treated) organoids. (B) Z-scores of the compounds labeled as toxic based on bright field image scoring, compared to Z-scores of dead organoids (treated for 24h with puromycin (3 mg/ml)) and viable organoids treated without FDA compound. Bars represent mean + SD, n=3 for toxic compounds, n=71 for dead and healthy controls.

### FDA-approved drugs increase FIS in homozygous W1282X/W1282X organoid lines

We next set out to identify FIS increasing compounds. A schematic of the workflow is shown in **Fig. 3A**. Four organoid lines homozygous for W1282X-CFTR were pretreated with VX-809 and VX-770 to increase baseline function of W1282X/W1282X, facilitating hit detection. Two FDA-approved compounds were combined in each well. F508del/S1251N organoids treated with VX-770 were used as positive control on each plate, allowing CV value and Z’-factor calculation for quality control purposes. Inter- and intra-assay CV values were 17% and 14% respectively, averaged over the four donors, highlighting robustness of the assay. For each donor two 384-wells plates were measured in triplicate, resulting in 6 Z’-factors for donor 1 and 5 Z’-factors for donors 2-4 due to technical errors. Average Z’-factors for all plates approximated 0.4, ranging between 0.3 and 0.5 **(Fig. 3B)**.

**Figure 3:**
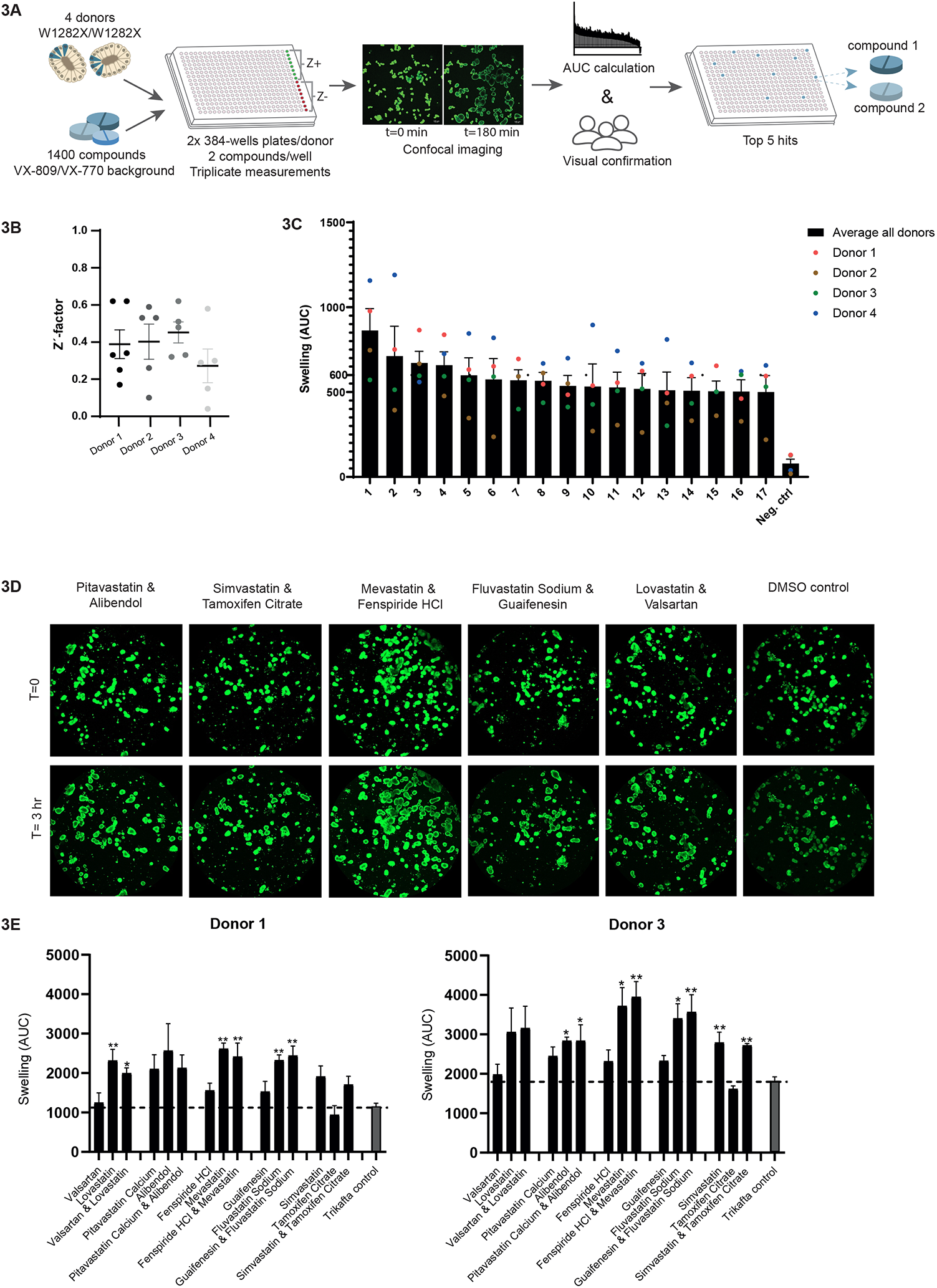
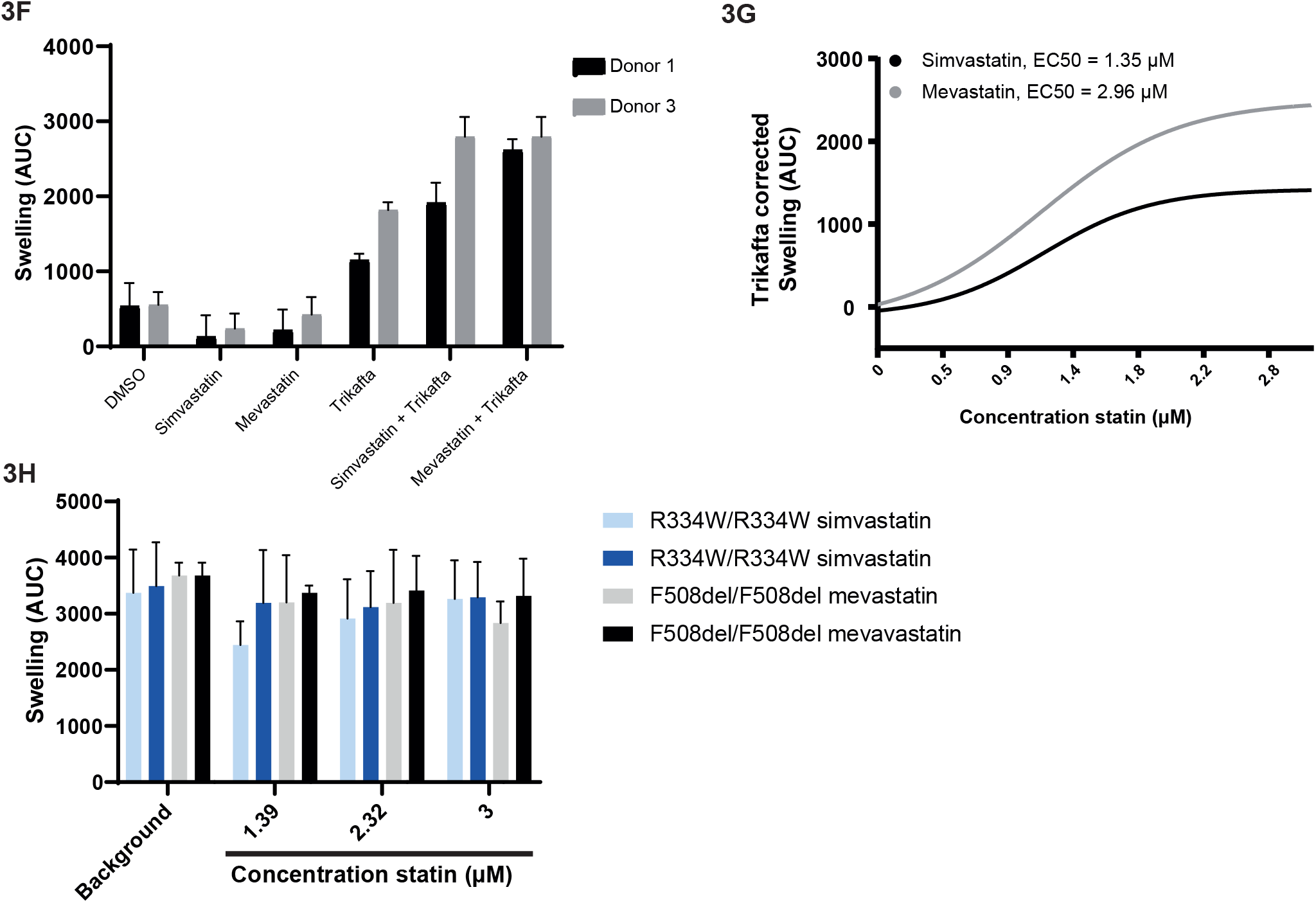
Schematic of workflow of FDA approved drug library testing on W1282X/W1282X PDIOs. (B) Z’-factor for each 384-wells plate for the four PDIOs. As positive control, F508del/S1251N PDIOs treated with VX-770 were used. (C) AUC values of the top 17 compound combinations, averages of the triplicate experiments of all PDIOs are shown with the different PDIOs indicated by the 4 different coloured dots. (D) Pictures of the PDIO line 1, of the primary screen, for the top 5 compound combinations and the DMSO control (t=0 and t=3 hours). (E) AUC values of the secondary FIS screen in 2 W1282X/W1282X PDIOs, in which compounds of the primary screen top 5 were tested separately and as original combination. Trikafta (CFTR modulators VX-445/VX-661/VX-770) was used as background instead of VX-770/VX-809. (F) AUC values for two W1282X/W1282X PDIOs upon treatment with the statins and CFTR modulators seperately (all at (3 µM), as well as the combination. (G) AUC values for one W1282X/W1282X PDIO upon a mevastatin and simvastatin concentration gradient, corrected for Trikafta induced baseline swelling. (H) AUC values for R334W/R334W and F508del/F508del PDIOs upon simvastatin and mevastatin treatment, with respectively VX-770 and VX-770/VX-809 as background (all at 3 µM).

The average AUC of the replicate experiments for all treatments are shown in **Sup. Figure 1**. 17 compound combinations resulted in an average increase of at least 500 AUC **(Fig. 3C)**. The top 5 compound combinations reaching the highest AUC values were verified by visual analysis **(Fig. 3D & Table 3)**. Subsequently, P-values were calculated for the difference between the average of each plate and the FDA-compound treated wells. Prior to multiple testing correction of the p-values by the Benjamin-Hochberg false discovery rate (FDR) method, 40 compound combinations resulted in a significant increase (p<0.05) in AUC compared to the average of the plate **(Supplemental Table 2)**. Included in this 40 compound combinations were the top 5 hits based on AUC values and 9 out-of-12 of the rest of the top 17. However, after FDR correction only the positive controls remained significantly different. Still, interestingly one of each of the compounds of the top 5 compound combinations is a statin, suggesting a statin-induced effect on organoid swelling. Altogether the results provided sufficient rationale to confirm whether these statins have indeed potential to increase CFTR function.

**Table 3:**
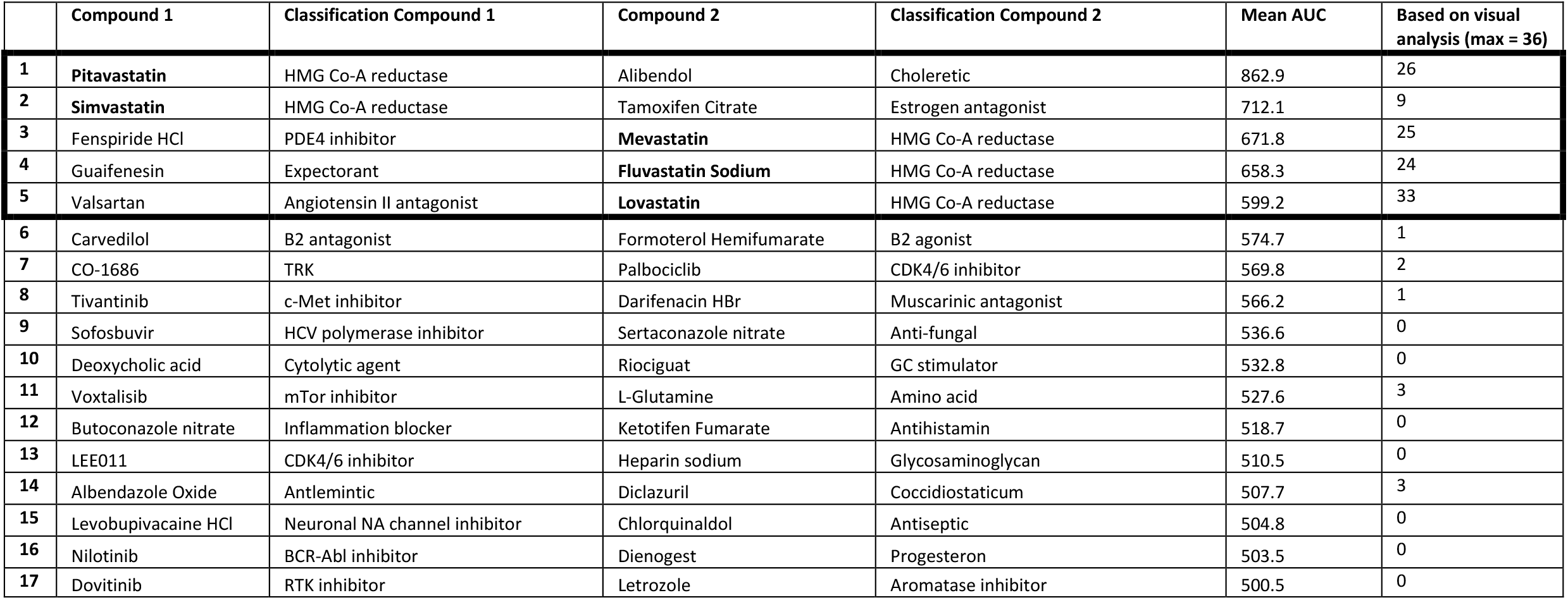
Hits based on mean AUC over all donors (>500)

To assess whether the statins of the compound combinations were responsible for AUC increase, the top five different compound combinations were repurchased from an independent supplier and tested on W1282X/W1282X donors 1 and 3. Compounds were tested in combination with VX-445/VX-661/VX-770, to further maximize the window of opportunity to detect effects in comparison to the primary screen **(Fig. 3E)**. The data of both donors combined confirmed that Simvastatin, Mevastatin, Fluvastatin and Lovastatin further induced organoid swelling, yet not significantly in both donors. Alibendol additionally resulted in a significant increase in AUC. In the absence of CFTR modulator treatment no Simvastatin or Mevastatin induced CFTR response was observed in both W1282X/W1282X donors **(Fig. 3F)**. By assessing swelling induction with a concentration range of Simvastatin and Mevastatin, we observed a concentration-dependent increase in AUC levels **(Fig. 3G)**. Concentrations above 3 µM resulted in decreased AUC values due to compound-induced toxicity (data not shown). Using 3 µM as maximal value, EC50s were respectively 1.35 and 2.96 µM for Simvastatin and Mevastatin, whereas a higher maximum swelling was induced with Mevastatin than with Simvastatin.

To evaluate whether the statins increased FIS of specifically W1282X CFTR, we assessed statin induced swelling in organoid cultures homozygously expressing F508del/F508del and R334W/R334W in combination with VX-809/VX-770 and VX-770, respectively. For these genotypes, we did not observe a further increase of FIS by Mevastatin and Simvastatin **(Fig. 3H)**.

## DISCUSSION

Whilst potent compounds have been developed for the most common F508del CFTR mutation, there’s a lack of medication for other genotypes and an urgent need to improve therapy for these CF patients. Approximately 10% of the worldwide CF population carry premature termination codons (PTC) mutations that result in production of truncated CFTR protein with severe loss-of-function. Developing new drugs is a time-consuming and expensive process. In this regard, drug repurposing is an attractive solution providing economical and time-wise benefits. In this study, we developed a 384-wells version of the forskolin induced swelling (FIS) assay, exemplifying the feasibility of large-scale compound screening with a functional read-out using patient-derived organoids.

We adapted several steps compared to the previously described 96-wells FIS assay ^12^. Importantly, medium replacement steps are not needed in this experimental pipeline, enabling a straightforward working scheme. To assure successful screening, quality and reliability of the 384-wells FIS assay was characterized. We show that replicate experiments yield similar results with low intra- and inter-assay CV values and without outer row or column associated effects. Overall, assay performance of the 384-wells FIS assay was adequate with Z’-factors reaching 0.4 or higher ^15^. Z’-factors were robust in the W1282X/W1282X screen as well, however, the positive control here consisted of F508del/S1251N organoids treated with VX-770 as a more suitable positive control specifically for W1282X/W1282X organoids was not available. Potentially in future studies the recently described RT agent DAP can be used for this ^22^ or a combination of compounds that together results in higher levels of CFTR rescue ^8^. In the future, additional automation such as automated organoid dispensers, drug printers and centrifugal washers might further reduce technical variability, thereby increasing the Z’-factor.

So far, mainly small-scale screenings using PDIOs have been performed, e.g. <100 compounds ^23,24^. Although recent work showed the implementation of organoids cultures in 3D matrix in a 384- and 1536-wells format for ultrahigh screening, the exploited read-outs, such as fluorescence-based viability analyses were relatively simple and exhibited a lower Z’-factor^25^. Jiang *et al*. (2021)^26^ show high-throughput therapy in the context of CFTR, using induced pluripotent stem cells (IPSCs) differentiated into early stage lung progenitor cells (LPCs) in a fluorescence-based assay of CFTR channel activity, yet with a lower Z’-factor (0.34). Berg *et al* (2019)^27^describe a 96-wells screening assay using 3D-cultured primary airway cells with a disease-relevant read-out consisting of liquid secretion quantification, yet Z’-factors were not described in this study. The clinical relevance of the exploited FIS assay in our study in combination with the robust Z’-factors in a 384-wells plate format, underline the strength of this study.

Whilst toxicity testing was not the main focus in our study, we exploited the 384-wells screening pipeline to analyze potential toxicity of the FDA-library. Indeed, 43 compounds induced toxicity and were excluded for further characterization. This approach could be translated to different PDIO systems, for example when comparing compound-induced toxicity in tumor and wildtype organoids derived from the same patient.

The effect of the 1400 FDA-approved compounds on CFTR mediated fluid secretion was assessed and PDIOs harbouring two PTC mutations, W1282X/W1282X, were exploited. As it has been shown that W1282X is the only CFTR PTC mutation that can to a minor extent be rescued by CFTR modulators, exploiting organoids with this genotype allows detection of a large range of hits encompassing several mode-of-actions (MoAs) ranging from nonsense-specific effect to more general CFTR modulating effects. 17 compound combinations resulted in an average increase of at least 500 AUC. Using a statistical approach, after multiple testing correction only the positive controls remained significantly different.

Altogether, the AUC analysis of the FIS experiments; the visual analysis to assess PDIO swelling in a binary way and the statistical analysis all pointed in the same direction concerning the top 5 compound combinations. Interestingly, one of each of the compounds of this top 5 compound combinations was a statin, suggesting a statin-induced effect on organoid swelling. Using newly purchased compounds from a different supplier, the top 5 compounds combinations were tested separately as well as original combination. CFTR modulators VX-770/VX-661/VX-445 were used as a background instead of VX-770/VX-809 due to its higher potency to increase CFTR function, thereby further maximizing the window of opportunity to detect effects, as well as their favorable safety profile and more clinically robust benefit ^28^. Indeed, with the exception of Pitavastatin, the four statins significantly increased AUC values of organoid swelling in comparison to the background alone.

In order to assess whether the statins increased CFTR function only in W1282X/W1282X organoid cultures, we assessed the effect of Simvastatin and Mevastatin on R334W/R334W and F508del/F508del organoid cultures, in combination with VX-770 and VX-809/VX-770 respectively. However, here no further increase in CFTR function was observed. This points into the direction that the effect is W1282X/W1282X or PTC specific. Interestingly, a HTS study using FRT cells stably expressing F508del-CFTR with a yellow fluorescent protein (YFP) flux assay as read-out, also revealed Atorvastatin calcium and Fluvastatin as hits ^29^. In future studies, assessing the effect of statins on a large panel of organoid with different genotypes will be valuable to draw firm conclusions on this.

Additionally, in combination with genotype specificity studies, more research is needed to elucidate MoA. Whilst statins are generally known for their potential to inhibit HMG-CoA reductase, thereby inhibiting the production of cholesterol and isoprenoids ^30,31^, their potential to increase CFTR function is in fact contradictory to previous studies. By contributing to the creation and maintenance of the lipid rafts in which CFTR resides, cholesterol has been described to positively impact CFTR levels and function ^32^. However, in this study more fundamental techniques were exploited and results were not verified in primary patient-derived cells. Statins additionally have been described to interfere with STAT1, p38 MAPK and Akt phosphorylation ^33^, pointing into the direction of many other potential MoAs. Along with statins, Alibendol resulted in elevated AUC levels. Alibendol is an antispasmodic, choleretic and cholekinetic drug that has not been described previously to restore CFTR function^34^. Future MoA studies can guide novel directions for drug development, and additionally the field of medical chemistry can aid in further increasing efficacy of these compounds.

In conclusion, we have miniaturized our FIS assay into a robust 384-wells plate high-throughput screen and used it to assess toxicity and CFTR increasing potential of an FDA approved drug library. We found that statins increased CFTR function in PTC harbouring CF PDIOs, a mechanism that has not been previously described. The developed pipeline in this study has significant value as proof-of-principle study, showing the potential of performing high throughput screening using PDIOs in an assay with a functional read-out.

## MATERIALS AND METHODS

### Collection of primary epithelial cells of CF patients

All experimentation using human tissues described herein was approved by the medical ethical committee at University Medical Center Utrecht (UMCU; TcBio#19-831). Informed consent for intestinal tissue collection, generation, storage, and use of the organoids was obtained from all participating patients. Biobanked intestinal organoids are stored and catalogued (https://huborganoids.nl/) at the foundation Hubrecht Organoid Technology (http://hub4organoids.eu) and can be requested at.

### Human intestinal organoid culture

Human intestinal organoid culture was executed as described by Vonk *et al* (2020)^12^.

### Compounds

The FDA library, purchased from SelleckChem (Z178323-100uL-L1300), was stored at -80°C. All other compounds described in this study were purchased at SelleckChem and dissolved in DMSO at a 20 mM concentration.

### 384-wells FIS assay validation

Differences between the 96-wells FIS assay as previously described by Vonk *et al* (2020)^12^ and the 384-wells FIS assay described in this study, are summarized in **Table 1** in the Results section. To characterize forskolin induced swelling (FIS) in a 384-wells format, a quality replicate experiment was performed. Three 384-wells plates were seeded with F508del/F508del organoids and three with F508del/S1251N organoids, plates were seeded after three different organoid culture periods up to twelve weeks. The F508del/S1251N organoids were submerged in 8 µL complete culture medium, the F508del/F508del organoids in 8 µL complete culture medium supplemented with 3 µM VX-809. 24 hours later, FIS measurements were assessed in the presence of VX-770 (3 µM) and low (0.008 µM) and high (5 µM) forskolin (fsk) concentrations, resulting in min signal values and max signal values. Organoid swelling was monitored for 60 minutes using a Zeiss LSM 710 confocal microscope. Total organoid surface area per well was quantified based on calcein green staining using Zen Blue Software and area under the curve over time was calculated as described by ^12^. Coefficient of variation (CV) values were calculated according to the following formula: % CV= (SD of means)/(mean of means)x100.

Minimum and maximum swelling enabled Z’-factor calculation of each 384-well plate according to the following formula: Z’-factor = 1-(3x(σp – σn)/(μp-μn)), where σp is the standard deviation of the max signal wells (n=128 per plate, + CFTR modulator(s) and 5 µM fsk), σn is the standard deviation of the min signal wells (n=128 per plate, + CFTR modulator(s) and 0.008 µM fsk), μp is the mean of the max signal wells, μn is the mean of the min signal wells.

### Toxicity screen using 384-wells plates

Organoids were plated into 384-wells plates and were incubated for 24h with 8 µL complete culture medium supplemented with a single FDA compound per well at 3 µM (1443 compounds in total, divided over five 384-wells plates, in triplicate (twice with F508del/S1251N organoids, once with F508del/F508del organoids). Bright field images were taken per well and organoid viability was scored in a binary way (live/apoptotic) by three blinded investigators. To assess cellular toxicity in a quantitative way, organoids were stimulated with calcein green (7 µM) for 30 minutes and propidium iodide (0.1 mg/mL) for 10 minutes prior to confocal imaging. Total organoid area per well was determined based on total calcein green staining (Zeiss, excitation at 488 nM) and amount of dead cells per well was determined by total area of PI staining (Zeiss, excitation at 564 nM) using Zen Image analysis software module (Zeiss). The ratio of total area calcein green and total area PI (=T-score) was calculated to correct for varying number of organoids between wells. To further correct for the varying organoid sizes among plates the Z-score was calculated between the compound-treated organoids and control DMSO-treated organoids per plate. The Z-score was determined according to the following formula: z-score = (x-μ)/σ, where x is the calcein/PI ratio of each single FDA compound, μ is the mean calcein/PI ratio of 16 control wells on each plate, and σ is the standard deviation of the calcein/PI ratio of the same 16 control wells on each plate. Z-scores of the single FDA compounds were compared to Z-scores of wells treated (n=71) with a toxic concentration of puromycin (24h, 3 mg/ml). Z-scores that were below Q1-(3xIQR) of all the DMSO treated wells (Z-score=-2.5) or Z-score that were above Q3+(3xIQR) of all puromycin treated wells (Z-score=14.8) were excluded.

### FDA-screen using 384-wells plates

Four W1282X/W1282X organoid lines were plated into 384-wells plates and were incubated for 24h with 8 µL complete culture medium supplemented with two FDA compounds per well (1400 compounds in total, divided over two 384-wells plates) as well as VX-809 at 3 µM. The day after plating, FIS measurements were assessed in the presence of VX-770 (3 µM) and 5 µM forskolin (fsk). No FDA compounds were added to 8 wells per plate as negative control (= min signal), and F508del/S1251N organoids were added to 8 wells per plate and treated with VX-770 to serve as positive control (=max signal). Experiments were performed in triplicate with organoids of different passages. Organoid swelling was monitored as previously described for 180 minutes using a Zeiss LSM 710 confocal microscope. Additionally, bright field images were taken per well and organoid morphology was scored in a binary way (no swelling/swelling) by three blinded investigators.

### Conventional 96-wells format FIS assay

96-wells FIS assays were executed as previously described by ^12^. FIS of F508del/F508del and F508del/S1251N organoids w/w/o CFTR modulator(s) and increasing concentration of fsk performed in 96-wells plates was compared to FIS under similar conditions performed in 384-wells plates. F508del/F508del organoids were treated with VX-809 (3 µM) for 24h prior to FIS assays. VX-770 (3 µM) was added simultaneously with fsk for 1h. Z’-factors were calculated based on the minimal swelling signals induced with 0.008 or 0.02 µM fsk + CFTR modulators and the maximum swelling signals induced with 5 µM fsk + CFTR modulators. For the statin experiments in W1282X/W1282X, F508del/F508del and R334W/R334W donors, compounds were added 24 hours prior to FIS measurements with different combinations of CFTR correctors (3 µM). Before FIS measurements, VX-770 (3 µM) and fsk (5 µM) were added and organoid swelling was monitored during 180 minutes.

### Statistics

Statistical analysis was performed using GraphPad Prism 8.0. Data on the graphs are presented as mean ± SEM, as experiments were performed in triplicate with three technical replicates per biological replicate. One-way analysis of variance (ANOVA) analyses with Dunnet’s posthoc test were performed to analyze the differences in the secondary W1282X/W1282X screen, where a separate test was performed for each compound combination group to compare compound A/compound B or compound combination AB to the Trikafta background. Differences were considered significant at a P-value of <0.05. Statistical analysis of the primary W1282X/W1282X screen was performed in RStudio, averages were calculated of the replicate experiments of the 4 donors combined for all compound combinations. To analyze whether these averages increased FIS in comparison to the average of the plate, one-way Students T-Tests were performed and p-values were adjusted for multiple testing using the Benjamin-Hochberg false discovery rate (FDR) method.

## Acknowledgements

We would like to thank the people with CF who gave informed consent for generating and testing their individual organoids; all members of the research teams of the Dutch CF clinics that contributed to this work; all colleagues of the HUB Organoid Technology for their help with generating intestinal organoids and we thank Prof. Dr. Rene Eijkemans for his valuable statistical advise and help.

## Author Contribution Statement

S.S. and E.d.P. contributed to the design of the study, the acquisition, verification, analysis and interpretation of the data and have drafted the manuscript. N.I., S.W.F.S., A.M.V., J.E.B., E.K. D.M. and M.C.H. contributed to the acquisition of study data and revised the manuscript. C.K.v.d.E and J.M.B. have made substantial contributions to the conception and design of the study, interpretation of data and revised the manuscript.

---

**Supplemental Figure 1:**
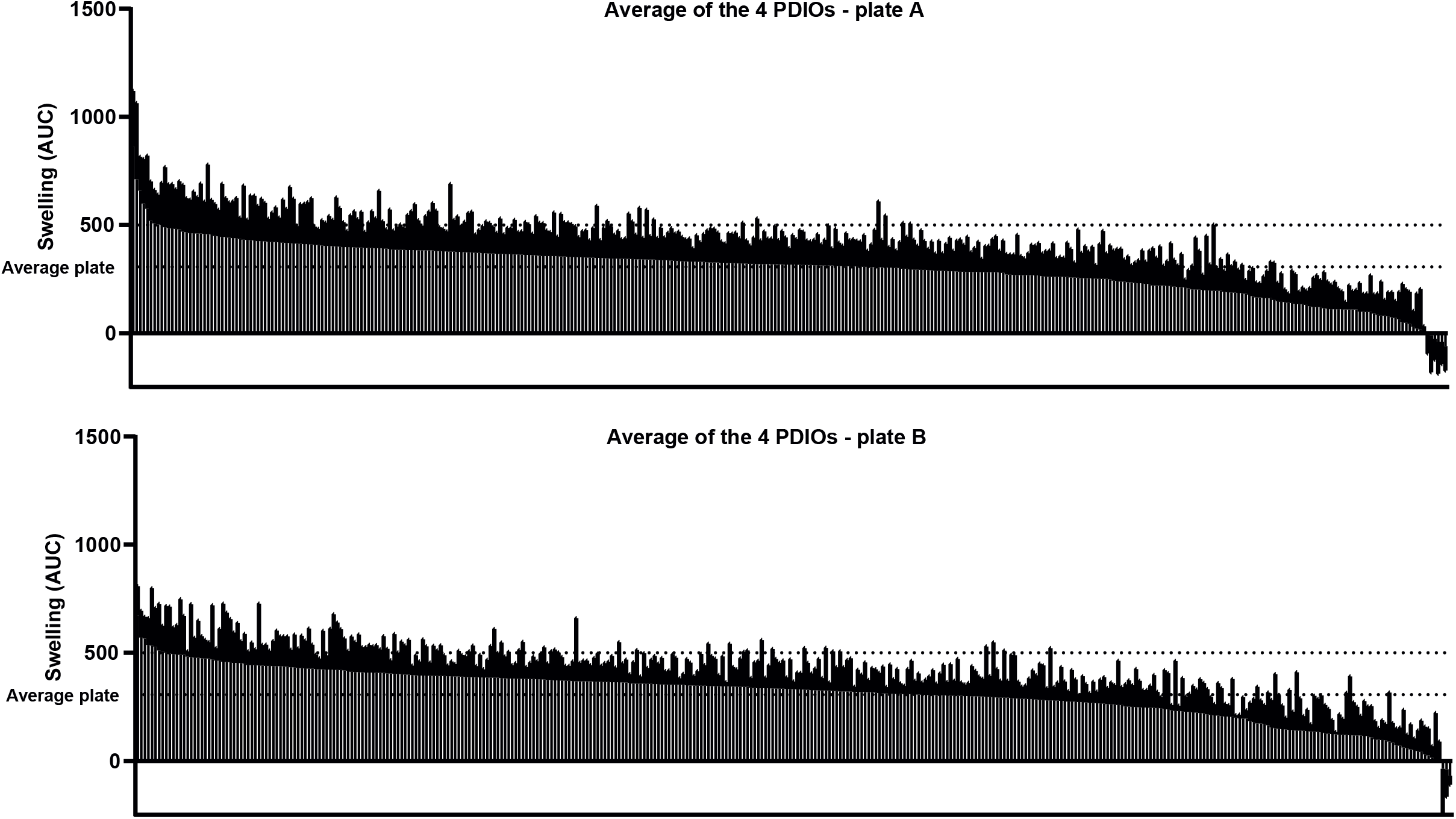
AUC values obtained in the FIS assay of the primary FDA-approved compound screen. Averages of the three replicates of the four W1282X/W1282X PDIOs are shown for each compound combination, for both 384-wells plates.

