## Supplemental Tables for "High-Throughput Functional Assay in Cystic Fibrosis Patient-Derived Organoids Allows Drug Repurposing"

**Supplementary Table 1. List of 43 compounds included for further screening, based on toxicity assay.**

| Compound number | Compound name | In vivo application |
|---|---|---|
| 1 | ABT-263 (Navitoclax) | Anti-cancer drug |
| 2 | YM155 (Sepantronium Bromide) | Anti-cancer drug |
| 3 | Bortezomib (PS-341) | Anti-cancer drug |
| 4 | Panobinostat (LBH589) | Anti-cancer drug |
| 5 | CEP-18770 (Delanzomib) | Anti-cancer drug |
| 6 | 17-AAG (Tanespimycin) | Anti-cancer drug |
| 7 | Ganetespib (STA-9090) | Anti-cancer drug |
| 8 | Saracatinib (AZD0530) | Anti-cancer drug |
| 9 | Onalespib (AT13387) | Anti-cancer drug |
| 10 | Dasatinib (BMS-354825) | Anti-cancer drug |
| 11 | Docetaxel (RP56976) | Anti-cancer drug |
| 12 | Ispinesib (SB-715992) | Anti-cancer drug |
| 13 | Paclitaxel (NSC 125973) | Anti-cancer drug |
| 14 | Rigosertib (ON-01910) | Anti-cancer drug |
| 15 | Epothilone B (EPO906, Patupilone) | Anti-cancer drug |
| 16 | Flavopiridol (Alvocidib) | Anti-cancer drug |
| 17 | Topotecan (NSC609699) HCl | Anti-cancer drug |
| 18 | Epirubicin (IMI 28) HCl | Anti-cancer drug |
| 19 | Tamoxifen (ICI 46474) | Anti-cancer drug |
| 20 | Vincristine (NSC-67574) | Anti-cancer drug |
| 21 | Rufinamide | Anti-epileptic/seizure drugs |
| 22 | Volasertib (BI 6727) | Anti-cancer drug |
| 23 | Neratinib (HKI-272) | Anti-cancer drug / Inhibits Xenograph growth |
| 24 | Ixazomib (MLN2238) | Anti-cancer drug |
| 25 | Crystal Violet | triarylmethane dye |
| 26 | Omipalisib (GSK2126458, GSK458) | Anti-cancer drug |
| 27 | Carfilzomib (PR-171) | Anti-cancer drug |
| 28 | Daunorubicin (RP 13057) HCl | Anti-cancer drug |
| 29 | Cabazitaxel (XRP6258) | Anti-cancer drug |
| 30 | Bisacodyl | Laxative drug |
| 31 | Dinaciclib (SCH727965) | Anti-cancer drug |
| 32 | Vinorelbine Tartrate | Anti-cancer drug |
| 33 | Digoxin | Atrial fibrillation and heart failure drug |
| 34 | Puromycin (CL13900) 2HCl | Aminonucleoside antibiotic |
| 35 | Vinblastine (NSC-49842) sulfate | Anti-cancer drug |
| 36 | Birinapant (TL32711) | Anti-cancer drug |
| 37 | Dasatinib Monohydrate | Anti-cancer drug |
| 38 | Docetaxel Trihydrate | Anti-cancer drug |
| 39 | Riociguat (BAY 63-2521) | Pulmonary hypertension drug |
| 40 | Mebendazole | Anti-cancer drug |
| 41 | Gemcitabine (LY-188011) HCl | Anti-cancer drug |
| 42 | Vancomycin HCl | Antibacterial agent |
| 43 | Fosbretabulin (Combretastatin A4 Phosphate (CA4P)) Disodium | Anti-cancer drug |

**Supplementary Table 2. List of p-values and adjusted p-valued (Benjamin-Hochberg method) of top 40 compound combinations in the W1282X/W1282X screen, based on a one-sided T-test where the average AUC value per condition over the 4 donors was compared to the average of the plate.**

| Compound_ID | p | p_adjusted | Compound 1 | Compound 2 |
|---|---|---|---|---|
| Pos. Control plate A | 0 | 0 | | |
| Pos. Control plate B | 0 | 0 | | |
| 1 | 0.000341 | 0.084206 | Fenspiride HCl (S4090) | Mevastatin (S4223) |
| 2 | 0.000683 | 0.126271 | Pitavastatin Calcium (S1759) | Alibendol (S1928) |
| 3 | 0.002352 | 0.348096 | Simvastatin | Tamoxifen Citrate (S1972) |
| 4 | 0.003027 | 0.365673 | CO-1686 (AVL-301) | Palbociclib (PD-0332991) HCl |
| 5 | 0.003459 | 0.365673 | Valsartan (S1894) | Lovastatin (S2061) |
| 6 | 0.008797 | 0.758291 | Guaifenesin (S1740) | Fluvastatin Sodium (S1909) |
| 7 | 0.010058 | 0.758291 | Nilotinib (AMN-107) | Dienogest |
| 8 | 0.010525 | 0.758291 | Carvedilol | Formoterol Hemifumarate |
| 9 | 0.016278 | 0.758291 | Butoconazole nitrate | Ketotifen Fumarate |
| 10 | 0.017213 | 0.758291 | Droperidol | Fluorometholone Acetate |
| 11 | 0.017866 | 0.758291 | Sotrastaurin | Ethynodiol diacetate |
| 12 | 0.018242 | 0.758291 | Linezolid | Acetylcysteine |
| 13 | 0.020269 | 0.758291 | S- (+)-Rolipram (S2127) | Naphazoline HCl (S2519) |
| 14 | 0.022949 | 0.758291 | Varlitinib | Entacapone |
| 15 | 0.0233 | 0.758291 | Cabazitaxel | Pergolide mesylate |
| 16 | 0.025466 | 0.758291 | Pravastatin sodium | Mirabegron |
| 17 | 0.025704 | 0.758291 | Voxtalisib (XL765, SAR245409) | L-Glutamine |
| 18 | 0.026123 | 0.758291 | Levobupivacaine HCl | Chlorquinaldol |
| 19 | 0.026898 | 0.758291 | Idasanutlin (RG-7388) | Lypressin Acetate |
| 20 | 0.031113 | 0.758291 | Cobicistat (GS-9350) | Eletriptan HBr |
| 21 | 0.031239 | 0.758291 | Triciribine | Cilostazol |
| 22 | 0.03282 | 0.758291 | Rosuvastatin Calcium | Secnidazole |
| 23 | 0.033328 | 0.758291 | Foretinib (GSK1363089) | Cilnidipine |
| 24 | 0.033605 | 0.758291 | TH-302 | Estradiol valerate |
| 25 | 0.035348 | 0.758291 | Cilengitide | Empty |
| 26 | 0.035555 | 0.758291 | Sofosbuvir (PSI-7977, GS-7977) | Sertaconazole nitrate |
| 27 | 0.037549 | 0.758291 | Probucol | Spectinomycin HCl |
| 28 | 0.038516 | 0.758291 | Marbofloxacin | Monobenzone |
| 29 | 0.041068 | 0.758291 | Griseofulvin | Florfenicol |
| 30 | 0.041323 | 0.758291 | Lomerizine HCl | Amoxapine |
| 31 | 0.041537 | 0.758291 | VX-680 (Tozasertib, MK-0457) | Celecoxib |
| 32 | 0.04302 | 0.758291 | Edoxaban | Nafarelin Acetate |
| 33 | 0.044306 | 0.758291 | Bendamustine HCl | Omeprazole |
| 34 | 0.046228 | 0.758291 | Licofelone | Sulfadoxine |
| 35 | 0.046428 | 0.758291 | Albendazole Oxide | Diclazuril |
| 36 | 0.047365 | 0.758291 | KPT-330 | Oxytocin (Syntocinon) |
| 37 | 0.048372 | 0.758291 | Retapamulin | Tiratricol |
| 38 | 0.049341 | 0.758291 | Sorafenib Tosylate | Rufinamide |
